# Mild weight loss promotes plaque growth and synthesis of pro-atherogenic lipids in ApoE-deficient mice fed a high-fat, high-cholesterol diet

**DOI:** 10.1101/2025.08.18.670993

**Authors:** Yun Zhu, Yanzhe Xu, Gerhard Liebisch, Jürgen Borlak

## Abstract

**Background:** According to the recent ACC/AHA Guideline on the Primary Prevention of Cardiovascular Disease, achieving an initial weight loss of ≥5% over six months in individuals with obesity is associated with clinically meaningful reductions in cardiovascular risk factors. However, emerging evidence suggests that weight loss may paradoxically increase the risk of CVD and atherosclerosis in certain populations. Motivated by these counterintuitive findings, we hypothesized that mild weight loss may induce metabolic reprogramming that promotes atherogenesis, potentially via an upregulation of pro-atherogenic lipids.

**Methods:** Apolipoprotein E-deficient mice were fed either a standard Chow or a high-cholesterol/high-fat (HCHF) diet for six months. A subset in each group underwent mild caloric restriction to achieve ∼5% body weight reduction. Atherosclerotic plaque burden in the brachiocephalic artery was assessed histologically. Hepatic lipidomic profiling was conducted via ESI–MS/MS, and lipid changes were compared with published plasma lipidomic data from obese human subjects stratified by CVD and diabetes status. Cardiovascular risk was further evaluated using the ceramide risk score (CERT1).

**Results:** Mild weight loss induced significant plaque growth, increasing lesion area by 28% in Chow-fed and 53.8% in HCHF-fed mice relative to controls. In HCHF-fed mice, caloric restriction markedly elevated free cholesterol and multiple pro-atherogenic phospholipids, including nine phosphatidylcholine, six phosphatidylethanolamine, one lysophosphatidylcholine, two sphingomyelin, and three ceramide species. Seven cholesterol esters also increased, and a CERT1 score of 8.1 indicated high CVD risk. These pro-atherogenic lipids were confirmed by comparison with published plasma lipidomic data from CVD and diabetes patients. In contrast, weight loss under Chow conditions caused minimal lipid alterations, with only isolated increases in phosphatidylcholine and lysophosphatidylcholine species.

**Conclusions:** Mild weight loss under high-fat dietary conditions accelerates atherosclerosis through increased production of pro-atherogenic lipids. These findings underscore the need to monitor lipidomic responses during weight loss interventions in obese individuals with CVD risk.

**Graphical Abstract:** 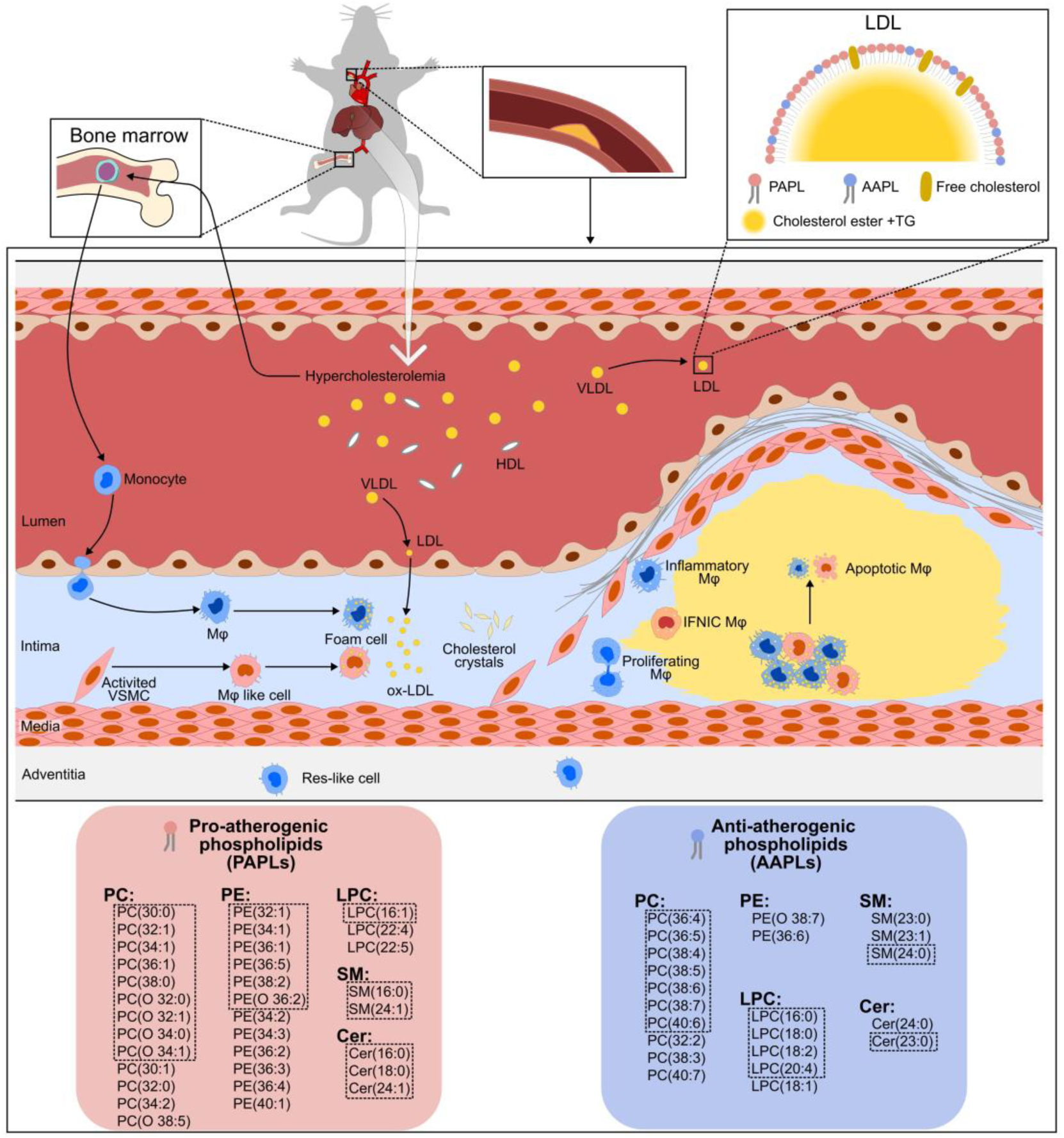

**Graphical Abstract model of phospholipid-driven atherosclerosis progression.**

Graphical summary depicting the mechanistic link between hypercholesterolemia, phospholipid composition, and atherosclerotic plaque development. The left panel highlights pro-atherogenic phospholipids, while the right panel shows anti-atherogenic phospholipids, both grouped by class and correlated with plaque size. Phospholipids enclosed within the dashed box represent those exhibiting statistically significant changes.

**Highlights:**

- Mild weight loss (∼5%) in ApoE-deficient mice fed a high-fat, high-cholesterol diet led to a paradoxical increase in atherosclerotic plaque burden.
- Weight loss under metabolically adverse dietary conditions may induce pro-atherogenic lipid remodelling that exacerbates atherosclerosis.
- Careful monitoring of atherogenic lipid species may be warranted in obese patients with existing cardiovascular disease undergoing dietary interventions.
- Comprehensive, tailored lifestyle interventions—beyond caloric restriction alone—may be necessary to avoid unintended cardiovascular harm during weight loss.

## Introduction

Weight loss is frequently recommended for individuals with cardiovascular disease (CVD), especially those who are overweight or obese, due to its association with improvements in metabolic risk factors such as insulin resistance ^1^. However, emerging evidence suggests that weight loss—particularly when unintentional or accompanied by weight fluctuations—may also be associated with adverse cardiovascular outcomes. For instance, in the ADVANCE trial of 10,081 adults with diabetes, both weight loss and gain were linked to an increased risk of cardiovascular events and all-cause mortality ^2^. Similarly, the ORIGIN trial, which included 12,521 individuals with prediabetes or type 2 diabetes, found that weight loss was independently associated with higher mortality, particularly among those with obesity ^3^. These findings raise concerns about the potential risks of weight variability in patients with cardiometabolic disease.

Weight cycling—defined as repeated episodes of weight loss followed by regain—is a particularly relevant concern in the context of GLP-1 receptor agonist therapy. The STEP 4 trial demonstrated significant weight regain after discontinuation of semaglutide ^4^. Furthermore, discontinuation of GLP-1 receptor agonists in patients with type 2 diabetes has been associated with increased risk of major cardiovascular events, regardless of prior CVD history ^5^. Supporting these observations, Bangalore et al. ^6^ reported that fluctuations in body weight were independently associated with increased mortality and cardiovascular events in a cohort of 9,509 individuals with established CVD.

While these associations remain the subject of debate—partly due to confounding factors such as unintentional weight loss from underlying illness and differences in baseline health status—they underscore the complex and incompletely understood relationship between weight dynamics and cardiovascular outcomes.

Further insight into this relationship comes from studies of subclinical atherosclerosis. The Canadian Longitudinal Study on Aging (CLSA) found that moderate weight loss was associated with increased carotid intima–media thickness (cIMT), a surrogate marker of atherosclerosis, in over 20,000 adults with baseline obesity ^7^. Similarly, the Coronary Artery Risk Development in Young Adults (CARDIA) study linked weight loss over time to increased coronary artery calcification, with cIMT remaining positively correlated with body mass index (BMI) regardless of the time point of measurement ^8^.

These findings align with the concept of the “obesity paradox,” in which overweight or mildly obese individuals with established CVD appear to have better survival outcomes— characterized by lower mortality and fewer cardiovascular events—than their normal-weight peers.

Together, these counterintuitive observations highlight the need for a mechanistic understanding of how body weight dynamics affect cardiovascular risk. To address this, we conducted a comprehensive lipidomic analysis in apolipoprotein E–deficient (ApoE⁻/⁻) mice, a well-established model of atherosclerosis. We hypothesized that mild weight loss may induce metabolic reprogramming that promotes atherogenesis via upregulation of pro-atherogenic lipid species, including cholesteryl esters, phospholipids, ceramides, and their acylated derivatives.

Given the liver’s central role in lipid metabolism and the synthesis of circulating atherogenic lipids, we focused our profiling on hepatic tissue. To assess translational relevance, we compared the hepatic lipid profiles from mice with plasma lipidomic data from obese human subjects, stratified by CVD and diabetes status. Clinically relevant lipid species were identified through lipidome-wide association studies linking specific lipids to cardiometabolic risk ^9,10^. This translational approach uncovered conserved pro-atherogenic lipids and suggests potential mechanistic pathways through which modest weight loss may paradoxically contribute to an increased atherosclerotic risk.

## Methods

### Ethics approval and consent to participate

Ethical approval was obtained from the local authority (Aktenzeichen: 50.203.2-INBIFO 11/02, Bezirksregierung Köln, Germany). As detailed in the original publication ^11^ all experimental procedures were conducted in accordance with institutional and international guidelines for the care and use of laboratory animals. Furthermore, the study adhered to the policies outlined by the American Association for Laboratory Animal Science (AALAS) for the humane care and use of laboratory animals.

### Animals

Male apolipoprotein E-deficient (ApoE⁻/⁻) mice (C57BL/6-apo Etm1Unc) were purchased from Charles River Laboratories (France). Animals were housed in polycarbonate cages with sterilized softwood bedding (Lignocel BK8/15, Altromin, Lage, Germany) under controlled environmental conditions, i.e. temperature (22 ± 2°C), relative humidity (55 ± 10%), and a 12-hour light/dark cycle. Throughout the study, animals were monitored for general health, appearance, and behaviour.

### Dietary intervention

To investigate the effects of long-term dietary modulation on hepatic lipid composition and bile acid remodelling (Supplementary Figure S1), ApoE-deficient mice were fed either a standard soybean-based diet containing 4.5% fat and 0.003% cholesterol, or an atherogenic high-cholesterol, high-fat (HCHF) diet consisting of 18.7% milk fat and 0.17% cholesterol (Supplementary Table S1 and S2), over a six-month period.

Additionally, to assess the impact of dietary restriction on the production of atherogenic phospholipids, a subset of ApoE-deficient mice underwent a mild 10% reduction in food intake.

Male ApoE-deficient mice aged 8–12 weeks were randomly assigned to the dietary groups. All diets were obtained from Sniff Spezialitäten GmbH (Soest, Germany). Unless otherwise specified, food and tap water were provided ad libitum.

Each experimental group included 10 animals. At the conclusion of the six-month intervention, liver tissues and bile samples were collected for comprehensive lipidomic and bile acid profiling.

Tissue and bile samples were provided by Dr. R. Schleef and are derived from sham animals from the original study by Holt et al. ^11^

### Plaque size in the brachiocephalic artery

Details of plaque size determinations have been described previously ^11^. Briefly, the brachiocephalic arteries were fixed in 4% formalin for 24 hours and subsequently embedded in paraffin. Serial cross-sections (5 μm thick) were cut and mounted onto glass slides. Sections were stained with hematoxylin and eosin as well as resorcin-fuchsin. Plaque size was quantified as the area occupied by atherosclerotic lesions within the vessel cross-section. For each mouse, five resorcin-fuchsin–stained sections, spaced 100 μm apart, were analysed with the using image analysis software (DISKUS®), and the values averaged. Images were captured and evaluated using a morphometric method ^12^. The relative plaque area was calculated by expressing the plaque area as a percentage of the total cross-sectional area of the vessel.

### Identification of Bile Acids and Their Conjugates

Bile acid profiling was performed using LC–ESI–MS/MS as previously described ^13–15^. Standards (>95% purity) were obtained from several suppliers. We analysed various bile acids and their conjugates, including tauro- and glyco-conjugates of cholic, chenodeoxycholic, deoxycholic, lithocholic, and ursodeoxycholic acids. Deuterated forms of selected bile acids served as internal standards (IS).

Quantification was based on the ratio of bile acid peak areas to their respective IS. For bile acids without identical deuterated IS, the closest IS by retention time and MRM transition was used.

### Lipid extraction of liver tissue and gall bladder bile

Liver samples were homogenized in phosphate-buffered saline (PBS) using an Ultra-Turrax homogenizer, with 300 µL PBS added per 500 mg of tissue. Five microliters of the resulting homogenate were used for lipid extraction according to the method of Bligh and Dyer ^16^, in the presence of non-naturally occurring lipid species serving as internal standards.

The following internal standards were included in the extraction: Cholesteryl esters (CE): CE 17:0, CE 22:0. Ceramides (Cer): Cer 18:1;O2/14:0, Cer 18:1;O2/17:0, Cer 18:1;O2[D7]/18:0. Free cholesterol (FC): FC[D7]. Glucosylceramides (GlcCer): GlcCer 18:1;O2/12:0, GlcCer 18:1;O2[D5]/18:0. Lyso-phosphatidylcholines (LPC): LPC 13:0, LPC 19:0. Phosphatidylcholines (PC): PC 14:0/14:0, PC 22:0/22:0. Phosphatidylethanolamines (PE): PE 14:0/14:0, PE 20:0/20:0 (di-phytanoyl). Phosphatidylserines (PS): PS 14:0/14:0, PS 20:0/20:0 (di-phytanoyl).

### Quantitative Lipidomics

We quantified lipids by direct flow injection electrospray ionization tandem mass spectrometry (ESI–MS/MS) in positive ion mode, using an established analytical setup ^17,18^. We employed specific precursor and neutral loss scans for lipid class identification and quantification: Phosphatidylcholine (PC), sphingomyelin (SM), and lyso-phosphatidylcholine (LPC) were detected using a precursor ion of m/z 184 ^17,19^. Phosphatidylethanolamine (PE) and phosphatidylserine (PS) were analysed via neutral losses of 141 and 185 Da, respectively^20^. PE-based plasmalogens (PE-pl) were analysed following the methodology described by Zemski-Berry ^21^. Ceramides (Cer) and hexosylceramides (HexCer) were detected via a fragment ion of m/z 264 ^22^. Free cholesterol (FC) and cholesteryl esters (CE) were quantified using a fragment ion of m/z 369, following selective derivatization of FC ^18^.

### Quantification of Cholesterol in Liver Tissue and Gallbladder Bile

Cholesterol and cholesterol esters were quantified using ESI–MS/MS in positive ion mode by comparison to analytical standards. Lipids were extracted from liver tissue and gallbladder bile and analyzed by direct flow injection using a validated bioanalytical assay ^18^. Free cholesterol (FC) and cholesterol esters (CE) were quantified by monitoring the fragment ion at m/z 369 Da following selective derivatization of FC, as described previously ^18^. Total cholesterol was calculated as the sum of FC and CE.

### Ceramide Risk Score (CERT1) Calculation

We adapted the scoring method from Laaksonen et al. ^23^ and calculated the ceramide risk score (CERT1) at baseline using tissue concentrations of four ceramide species: Cer(16:0), Cer(18:0), Cer(24:1), and Cer(24:0). For each animal, the absolute concentrations of Cer(16:0), Cer(18:0), and Cer(24:1), as well as their ratios to Cer(24:0), were evaluated against the distribution of all animals.

Scoring was assigned based on quartile distribution: Values in the 3rd quartile received 1 point, and those in the 4th quartile received 2 points. Cer(24:0) itself was not scored. The total CERT1 score thus ranged from 0 to 12. The following scale was used to assign cardiovascular risk categories based on the total score as follows: Low risk: 0–2; moderate risk: 3–6, increased risk: 7–9 and high risk: 10–12.

### Publicly available gene expression datasets

We retrieved gene expression datasets GSE190421 ^24^ and GSE28125 from the Gene Expression Omnibus (GEO; https://www.ncbi.nlm.nih.gov/geo/). Specifically, GSE190421 compares liver transcriptomes of ApoE deficient mice fed either a chow or high-fat diet (HFD), while GSE28125 contrasts liver gene expression profiles between wild-type and ApoE deficient mice under HFD conditions.

### Data Normalization and Statistical Analysis

For the dataset GSE28125, raw data pre-processing and normalization were performed using the Affy package (version 1.86.0) in R (version 4.5.0), applying the Robust Multi-array Average (RMA) method. Differentially expressed genes (DEGs) were identified using the Limma package.

For the RNA-sequencing datasets (GSE190421), we used the edgeR package (version 4.6.2) for an identification of DEGs. We applied the Benjamini–Hochberg (BH) procedure to control the false discovery rate (FDR) for multiple testing correction. Genes exhibiting a fold change (FC) ≥ 1.5 and an BH-adjusted p-value < 0.05 were considered significant.

### Statistical analysis

We performed power analysis in R using the pwr package to estimate the required group size for detecting statistically significant differences between the two dietary intervention groups, assuming a significance level (α) of 0.05 and a power (1−β) of 0.8. Based on total cholesterol, the estimated sample size was 9 animals. Therefore, a group size of 10 animals deemed sufficient for all planned comparisons.

Data distribution was assessed using the Shapiro–Wilk test for normality. For datasets not meeting normality assumptions, non-parametric tests including the Kruskal–Wallis test and the Wilcoxon rank-sum test (Mann–Whitney U test) were applied. For normally distributed data, we considered one-way analysis of variance (ANOVA) and unpaired two-tailed t-tests as appropriate.

Statistical analysis of plaque size in the brachiocephalic artery was performed with the GraphPad Prism software (version 9.0).

## Results

### Experimental Design and Body Weight Outcomes

Supplementary Figure S1 illustrates the overall experimental design, and Figure 1A shows the body weight changes of ApoE-deficient mice after 6 months on a Chow or high-cholesterol, high-fat (HCHF) diet. Mice fed the HCHF diet exhibited a 15.2% increase in body weight when compared to chow-fed controls. Dietary restriction (10% reduction in food intake) resulted in moderate decreases in body weight, i.e. a 6.7% reduction in the chow-restricted group (Chow_DR) and a 4.6% reduction in the HCHF-restricted group (HCHF_DR). However, the difference in body weight between the HCHF and HCHF_DR groups was not statistically significant. Overall, mild dietary restriction led to a small but statistically significant reductions in body weight.

**Figure 1.**
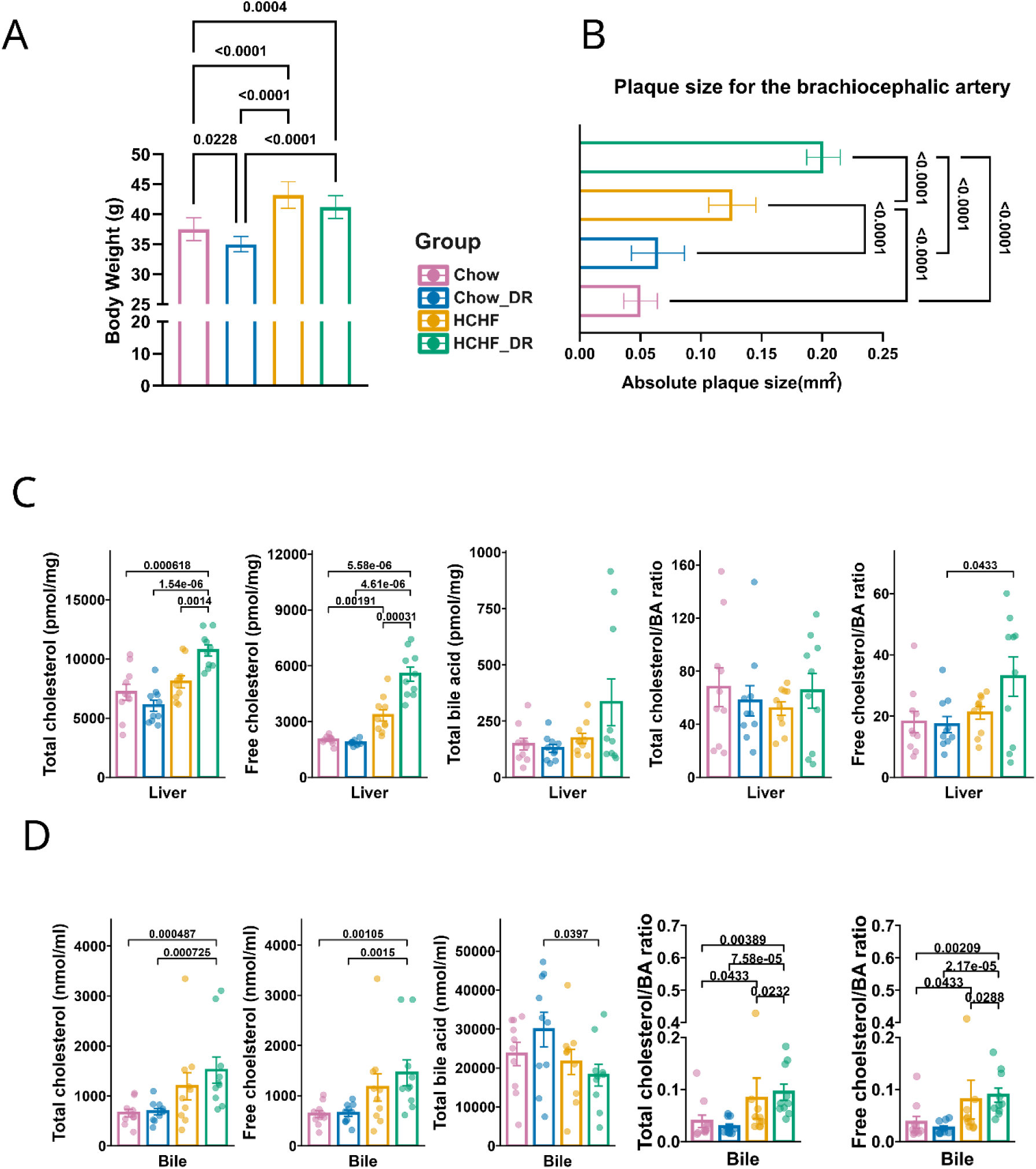
Body weight, atherosclerotic plaque size, and hepatic/biliary cholesterol and bile acid metabolism. (A) Body weight comparisons across the four dietary groups. (B) Morphometric analysis of plaque size in the brachiocephalic artery. (C) Hepatic levels of total cholesterol, free cholesterol, bile acids, and their ratios. (D) Biliary levels of cholesterol and bile acids with corresponding ratios. Data are presented as mean ± SEM. P values are shown where applicable.

### Atherosclerotic Plaque Development

Next, we assessed atherosclerotic plaque area in the brachiocephalic artery (Figure 1B). Chow-fed mice had an average plaque size of 0.05 mm², while mice on the HCHF diet showed a significantly larger average plaque area of 0.13 mm², representing a 2.6-fold increase (p < 0.001).

Interestingly, mild dietary restriction produced divergent effects depending on the dietary background. In the Chow_DR group, plaque area increased by 28% as compared to the unrestricted chow group, though this difference was not statistically significant. In contrast, the HCHF_DR group exhibited a 53.8% increase in plaque size relative to the unrestricted HCHF group, which was highly significant (p < 0.01). Finally, a comparison between the Chow and HCHF_DR groups revealed a 4-fold increase in plaque size, highlighting the exacerbating effect of HCHF feeding, especially following mild dietary restriction.

### Cholesterol homeostasis

Bar charts in Figure 1C and 1D present quantitative analyses of total cholesterol, free cholesterol, total bile acids (BA), and the cholesterol/BA ratio in liver tissue and gallbladder bile.

Mild dietary restriction exerted distinct effects on hepatic total cholesterol depending on the dietary background. Specifically, the Chow_DR group showed a non-significant 15.6% reduction in total hepatic cholesterol compared to ad libitum chow-fed controls. Conversely, mice fed the HCHF diet exhibited a 12.2% increase in total hepatic cholesterol (2.78 mg/g versus 3.12 mg/g liver weight). Although total cholesterol levels did not differ significantly between the ad libitum Chow and HCHF groups, free cholesterol was significantly elevated, i.e. 65.5% in HCHF-fed mice (p = 0.00191). The data are strikingly similar to human liver with increases in free cholesterol of 37.7% and 70.6% cases, respectively in MAFL and MASH cases ^25^. Extraordinarily, mild food restriction in the HCHF_DR group caused a further significant increase in free cholesterol, surpassing levels observed in the unrestricted HCHF group (p = 0.00031). These results suggest that dietary restriction modulates cholesterol metabolism in a diet-dependent manner, with free cholesterol increasing nearly threefold in the HCHF_DR group relative to Chow-fed controls.

The increase in free cholesterol is likely due to the marked repression of ACAT1 and over tenfold suppression of Ces1f, which likely contributed to cholesterol ester hydrolysis. Since hepatic lipase levels were unchanged, these changes together provide a plausible mechanism for the rise in free cholesterol despite stable total cholesterol between the two dietary groups (Figure 1C).

Considering the differences in cholesterol content between the diets and the role of excess cholesterol in bile acid (BA) synthesis, it was unexpected that the total cholesterol-to-BA ratio remained unchanged across dietary groups (Figure 1C). However, when focusing on free cholesterol, the cholesterol-to-BA ratio was significantly elevated in the HCHF_DR group, suggesting a potential impairment in bile acid synthesis.

We further analyzed the lipid composition of gallbladder bile, as summarized in Figure 1D. Total biliary cholesterol did not differ between Chow and Chow_DR groups. In contrast, mice fed the HCHF diet exhibited nearly a two-fold increase in total biliary cholesterol. Mild food restriction in HCHF-fed mice led to a further 27% increase in total cholesterol, although this did not reach statistical significance. Interestingly, despite the fact that the HCHF diet contains 56 times more cholesterol than the standard Chow diet, biliary total cholesterol increased by only two-fold, suggesting rapid saturation of cholesterol absorption or hepatic processing following HCHF intake.

A similar trend was observed for free cholesterol: levels were comparable between Chow-fed groups, whereas the HCHF group exhibited a borderline significant, nearly two-fold increase (p = 0.063) when compared to Chow and Chow_DR mice. Mild dietary restriction in the HCHF_DR group led to a further 24% increase in free cholesterol, which likewise did not reach statistical significance.

Although excess cholesterol is typically is converted into bile acids, total bile acid concentrations were paradoxically reduced in HCHF-fed groups. Specifically, the HCHF group exhibited a non-significant 10% decrease, while the HCHF_DR group showed a non-significant 23% reduction. Consequently, the cholesterol-to-BA ratios were significantly elevated in HCHF-fed mice, whether calculated using total or free cholesterol, with most pronounced differences observed between Chow_DR and HCHF_DR animals.

### Identification of clinically relevant atherogenic cholesterol esters

Previous research established a strong association between specific plasma cholesterol ester (CE) species and cardiometabolic risk in a cohort of 551 CVD and 775 T2DM patients ^9,10^. Building on this foundation, we analyzed the composition of cholesterol ester species with particular emphasis on clinically relevant atherogenic variants. The results for the liver are shown in Figure 2A (left panel). Although total CE levels did not differ significantly across dietary groups, marked differences emerged when individual CE species were examined.

**Figure 2.**
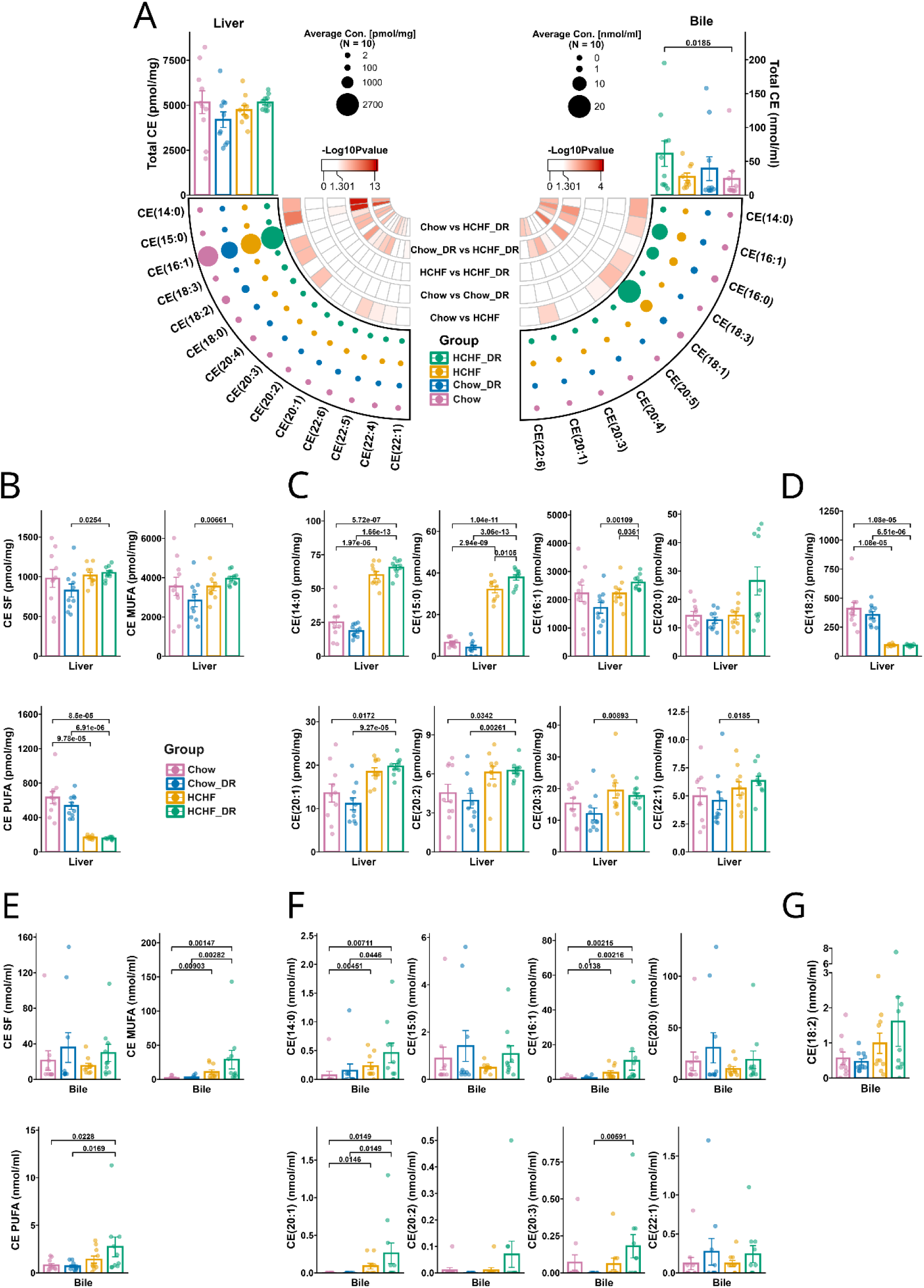
Cholesteryl ester (CE) profiling in liver and bile. (A) Circular bubble heatmaps illustrating the composition and concentrations (pmol/mg tissue) of CE species in the liver (left panel) and bile (right panel). Bubble size indicates average concentration (pmol/mg in liver or nmol/ml in bile); red arcs denote -log₁₀ transformed P values for group comparisons. (B) Liver CE subclass distribution, including saturated fatty acid (SFA), monounsaturated fatty acid (MUFA), and polyunsaturated fatty acid (PUFA) species. (C) Quantification of pro-atherogenic CE species in the liver. (D) Quantification of anti-atherogenic CE species in the liver. (E) CE class and subclass (SFA, MUFA, PUFA) concentrations in bile. (F) Quantification of pro-atherogenic CE species in bile. (G) Quantification of anti-atherogenic CE species in bile. Data are presented as mean ± SEM. P values are shown where applicable.

Specifically, cholesterol esters bound to saturated fatty acids (SFAs) and monounsaturated fatty acids (MUFAs) were significantly elevated in Chow_DR compared to HCHF_DR animals. Conversely, polyunsaturated fatty acid (PUFA)-bound cholesterol esters were significantly reduced in animals fed the HCHF diet, an effect that remained unchanged by mild food restriction (Figure 2B).

Several atherogenic CE species were increased in HCHF-fed animals, including CE(14:0), CE(15:0), CE(16:1), CE(20:1), CE(20:2), CE(20:3), and CE(22:1). Mild food restriction further augmented increases in certain atherogenic CE species, particularly CE(15:0) and CE(16:1), with CE(20:0) showing borderline significance (p = 0.1). Regarding anti-atherogenic cholesterol esters, CE(18:2) was significantly suppressed following the HCHF diet, and was unaffected by mild food restriction (Figure 2D).

The results for bile are presented in Figure 2A (right panel). Mild food restriction caused non-significant increases in SF-bound CE. Nonetheless, MUFA- and PUFA-bound CEs were significantly increased following the HCHF diet (Figure 2E). Pro-atherogenic CE species were significantly enriched in HCHF-fed animals and included CE(14:0), CE(16:1), CE(20:1), and CE(20:3). Mild food restriction led to further increases in CE(16:0), CE(20:0), CE(20:2) and CE(22:1), although these did not reach statistical significance (Figure 2F). Unexpectedly, the anti-atherogenic CE(18:2) was insignificantly upregulated in HCHF-fed animals (Figure 2G).

Collectively, these findings indicate a disruption in cholesterol homeostasis in response to the HCHF diet, which is further exacerbated by mild dietary restriction.

### Modulation of phospholipid classes following dietary interventions

Changes in hepatobiliary phospholipid composition in response to Chow and HCHF diets are shown in Figure 3. Compared to chow-fed controls, HCHF-fed animals exhibited a significant reduction in total hepatic phospholipids (p < 0.000236). However, mild dietary restriction in HCHF_DR animals significantly restored total phospholipid levels, bringing them close to those observed in chow-fed controls (Figure 3A1). In contrast, dietary restriction of chow-fed animals (Chow_DR) had no significant effect on total phospholipid content, indicating that the restorative effect of dietary restriction was specific to the HCHF dietary context.

**Figure 3.**
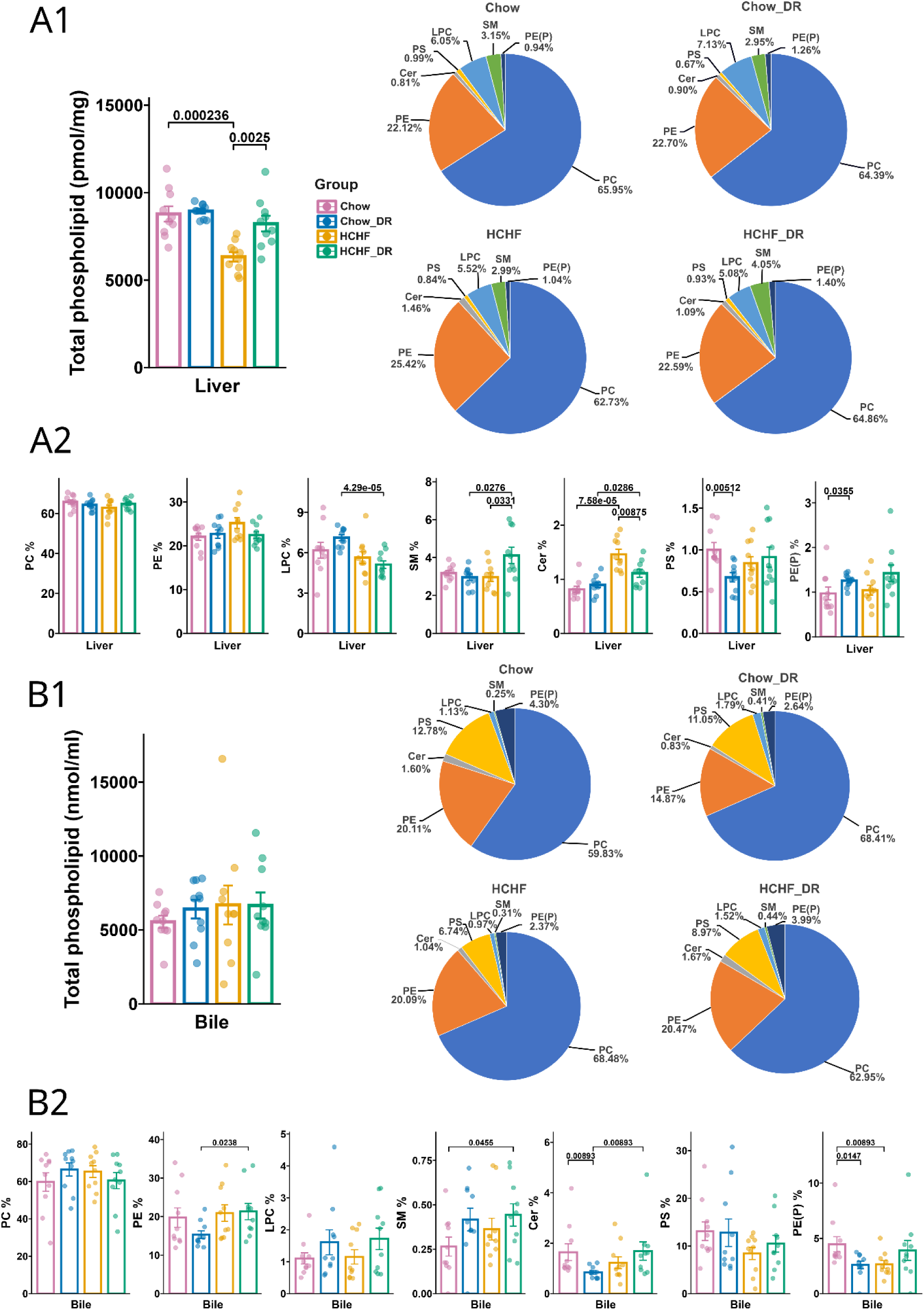
Total phospholipid content and phospholipid class composition. (A1–A2) Liver: total phospholipid concentration and class percentages (PC, PE, LPC, SM, Cer, PS, PE(P)). (B1–B2) Bile: phospholipid quantification and class distribution. Pie charts display average class composition. Data are presented as mean ± SEM. P values are shown where applicable.

The relative proportions of phospholipids in liver extracts are illustrated as pie charts in Figure 3A1, with corresponding statistical analyses presented in Figure 3A2. Among the major phospholipid classes, phosphatidylcholine (PC) and phosphatidylethanolamine (PE) levels remained consistent across all dietary groups (Figure 3A2). In contrast, lysophospholipids were significantly reduced in mice on the HCHF diet, particularly when comparing mildly food-restricted animals on the Chow versus HCHF regimens.

Note sphingomyelin and ceramide levels increased significantly in mice fed the HCHF diet— by 35% and 82%, respectively. These increases were further amplified under conditions of mild dietary restriction. Furthermore, in the Chow_DR group, phosphatidylserine (PS) levels were significantly decreased, while PE-based plasmalogens (PE(P)) were significantly elevated when compared to unrestricted Chow-fed animals.

We also considered the relative proportions of phospholipids in gallbladder bile which are shown in Figure 3B1 (pie charts), with corresponding significance values in Figure 3B2 (histograms). Total phospholipid and PC levels remained stable across all dietary groups. Although PE levels in the Chow_DR group showed a reduction from 20.11% to 14.87%, this change did not reach statistical significance (p = 0.127). Given the lack of difference in PE between Chow and HCHF diets, the statistical significance between Chow_DR and HCHF_DR is likely incidental. Furthermore, there were no significant changes in the proportions of lysophosphatidylcholine (LPC) or phosphatidylserine (PS) across any dietary group. However, gallbladder sphingomyelin levels increased in response to food restriction, achieving statistical significance only in the comparisons between Chow and HCHF_DR animals. Interestingly, ceramide levels were differentially regulated by food restriction, that is reduced by 52% in Chow-fed mice but increased by 60% in the HCHF_DR group. Lastly, PE-based plasmalogens were significantly reduced by food restriction in both Chow and HCHF-fed animals.

Together, the findings highlight the distinct and diet-dependent remodelling of hepatic and biliary phospholipid profiles in ApoE-deficient mice, emphasizing the complex interplay between dietary composition and mild food restriction on lipid metabolism.

### Phospholipidomics, identification and quantification of lipid species

Figure 4A provides a detailed overview of the relative distribution and quantification of phosphatidylcholine (PC) fatty acyl chains, with quantification indicated by the size of the bubbles. Only acyl chains that show statistically significant differences in at least one comparison across the dietary groups are included. The corresponding p-values are displayed in the inner circle, where the intensity of the red colour reflects the –log_10_ p-value, ranging from p < 0.05 to p < 10^−11^. For clarity, dietary interventions are color-coded: Chow and Chow_DR animals are represented in pink and blue, respectively, while animals on a HCHF and HCHF_DR diet are shown in yellow and green.

**Figure 4.**
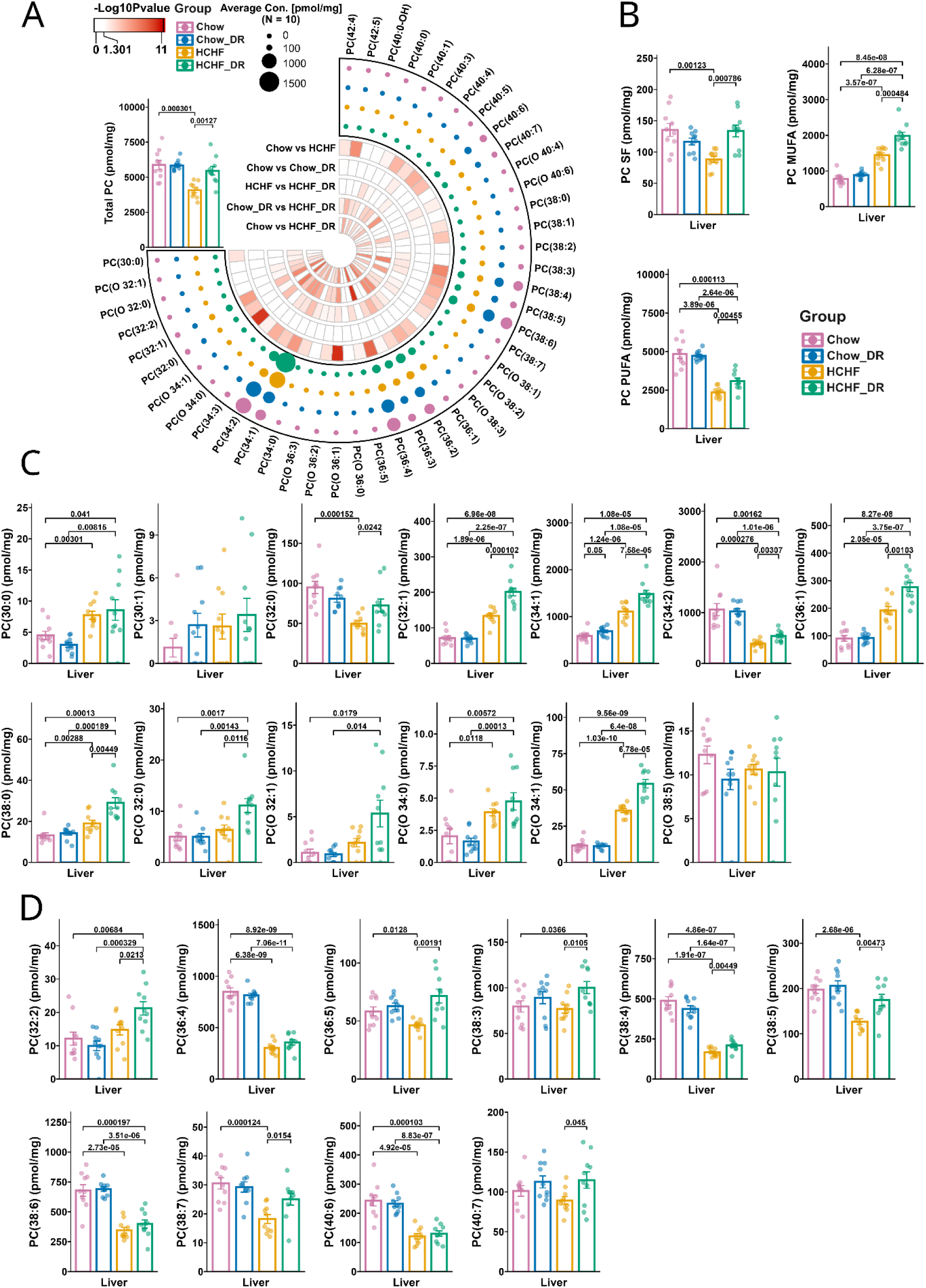
Phosphatidylcholine (PC) species composition in liver. (A) Circular bubble heatmap displaying the distribution and concentrations (pmol/mg tissue) hepatic PC species across diet groups. Bubble size represents average concentration (pmol/mg); red arcs indicate -log₁₀-transformed P values from group comparisons. (B) Total hepatic levels of PC species containing saturated fatty acids (SFAs), monounsaturated fatty acids (MUFAs), and polyunsaturated fatty acids (PUFAs). (C) Quantification of pro-atherogenic PC species in the liver. (D) Quantification of anti-atherogenic PC species in the liver. Data are presented as mean ± SEM. P values are shown where applicable.

Note total phosphatidylcholine (PC) content (Figure 4A) did not differ between Chow and Chow_DR animals. However, PC levels were significantly reduced in the livers of HCHF-fed mice compared to Chow-fed controls. While mild food restriction had no effect on hepatic PC levels in Chow-fed animals, it restored PC content in HCHF_DR mice to levels comparable to those observed in the Chow group.

Figure 4B depicts the distribution of fatty acyl chain classes within phosphatidylcholine (PC). Although the HCHF diet is significantly enriched in saturated fatty acids (SFAs), we observed a pronounced decrease in PC-bound saturated acyl chains in HCHF-fed animals. However, mild food restriction restored PC-bound SFA levels in HCHF_DR mice to those comparable with Chow-fed controls. In contrast, monounsaturated fatty acids (MUFAs) esterified to PC were significantly elevated in HCHF-fed animals and increased further with mild food restriction. These results indicate a diet-induced remodelling of PC composition, shifting from saturated toward monounsaturated fatty acids—a shift that is enhanced by mild food restriction. Importantly, stearoyl-CoA desaturase (SCD), the enzyme responsible for converting saturated fatty acids into MUFAs, was significantly upregulated in ApoE-deficient mice on a high-fat diet when compared to Chow-fed ApoE controls (see Supplementary Figure S2).

Regarding PC-bound polyunsaturated fatty acids (PUFAs), and given that the HCHF diet is milk fat-based, it was logic to observe an approximate 50% reduction in PC-PUFA levels following the HCHF diet, and a 36% reduction with the HCHF_DR diet.

### Quantification of clinically relevant pro-atherogenic PC acyl chains

Independent research established a strong association between the lipid composition in the liver, adipose tissue and plasma in MASLD and severe obese patients ^26^. Building on this framework, we focused on specific PC-bound acyl chains previously linked to pro-atherogenic effects. To strengthen our analysis, we employed a consensus approach integrating data from four independent studies ^27–30^, selecting PC lipid species with well-characterized atherogenic potential across nearly 500 CVD patients.

As shown in Figure 4C, we observed significant increases in several atherogenic PC species— including PC(30:0), PC(32:1), PC(34:1), PC(36:1), PC(38:0), PC(O 32:0), PC(O 32:1), PC(O 34:0), and PC(O 34:1)—in the livers of ApoE-deficient mice on a HCHF diet. Strikingly, mild food restriction further elevated levels of PC(32:1), PC(34:1), PC(36:1), PC(38:0), PC(O 32:0), PC(O 32:1), and PC(O 34:1); thus underscoring the paradoxical and potentially detrimental impact of mild dietary restriction on the accumulation of pro-atherogenic PC lipid species.

On the other hand, certain plasma phospholipids have been associated with favorable outcomes in coronary heart disease (CHD) and coronary artery disease (CAD) ^27^. Based on this, we investigated the regulation of phospholipid species known for their protective and anti-atherogenic properties. As shown in Figure 5D, several phosphatidylcholine (PC) species—specifically PC(36:4), PC(36:5), PC(38:4), PC(38:5), PC(38:6), PC(38:7), and PC(40:6)—were significantly decreased in the livers of ApoE-deficient mice fed a HCHF diet. This reduction highlights the detrimental effect of the HCHF diet on hepatic phospholipid species linked to cardiovascular protection. Interestingly, mild food restriction in the context of the HCHF diet led to significant increases in PC(32:2) and PC(38:3) by 76% and 25%, respectively.

**Figure 5.**
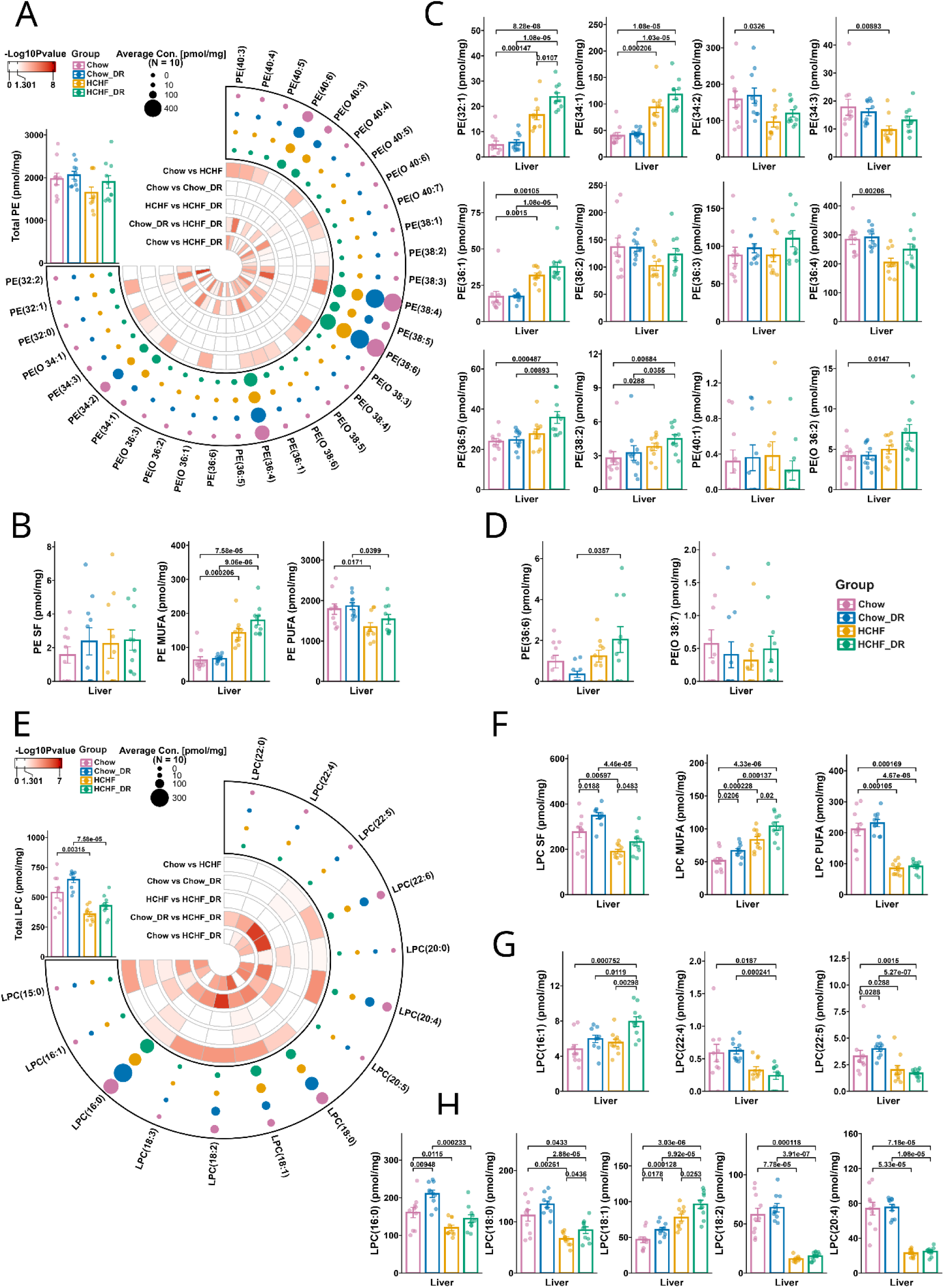
Quantification of hepatcic phosphatidylethanolamine (PE) and lysophosphatidylcholine (LPC) species. (A) Circular bubble heatmap displaying the concentrations (pmol/mg tissue) of PE species in the liver across different dietary groups. Bubble size indicates average concentration (pmol/mg); red arcs denote -log₁₀ transformed P values from one-way ANOVA. (B) Total hepatic phosphatidylethanolamine levels of saturated (SFA), monounsaturated (MUFA), and polyunsaturated (PUFA) PE species. (C) Quantification of pro-atherogenic PE species in the liver. (D) Quantification of anti-atherogenic PE species in the liver. (E) Circular bubble heatmap displaying the concentrations (pmol/mg tissue) of LPC species in the liver across different dietary groups. Bubble size indicates average concentration (pmol/mg); red arcs denote -log₁₀ transformed P values from one-way ANOVA. (F) Total hepatic lysophosphatidylcholine levels of saturated (SFA), monounsaturated (MUFA), and polyunsaturated (PUFA) LPC species. (G) Quantification of pro-atherogenic LPC species in the liver. (H) Quantification of anti-atherogenic LPC species in the liver. Data are presented as mean ± SEM. P values are shown where applicable.

We also quantified PC species in gallbladder bile, and the data are presented in supplementary Figure S3. Consistent with hepatic findings, PC species bound to SFAs and MUFAs were significantly elevated, while those bound to PUFAs were markedly reduced. Note several pro-atherogenic PC species—including PC(30:0), PC(32:1), PC(34:1), PC(36:1), PC(38:0), PC(O-34:1), PC(O-34:0), and PC(O-38:5)—were significantly increased in the bile of ApoE-deficient mice on the HCHF diet. Mild food restriction further exacerbated the elevation of PC(34:1).

With regard to protective and anti-atherogenic PC species, we observed significant increases in PC(32:2), PC(38:3), and PC(40:7), whereas levels of PC(36:4) and PC(38:4) were reduced.

Overall, the HCHF diet induced remodelling of PC composition. Although total PC content was significantly reduced, the diet promoted the enrichment of pro-atherogenic PC acyl chains.

### Quantification of clinically relevant pro-atherogenic PE and LPC acyl chains

Phosphatidylethanolamine (PE) is the second most abundant phospholipid species in the liver (Figure 5A), and following a HCHF diet, total hepatic PE levels were reduced by 16%, although this decrease did not reach statistical significance (p = 0.1). In contrast, mild food restriction restored PE levels to those observed in chow-fed controls.

Further analysis of PE fatty acid composition revealed no significant differences in SFA-bound PE species across dietary groups (Figure 5B). However, MUFA-bound PE were significantly elevated in HCHF-fed mice and increased further following mild dietary restriction. Conversely, PUFA-bound PE species were significantly reduced in both HCHF and HCHF_DR groups. These data suggest a diet-induced remodelling of PE composition, characterized by a shift from PUFA-enriched to MUFA-enriched species. This shift was modestly, though not significantly, enhanced by mild food restriction (p = 0.0618).

Next, we quantified atherogenic phosphatidylethanolamine (PE) species in the livers of ApoE-deficient mice. Several species—PE(32:1), PE(34:1), PE(36:1), PE(36:5), PE(38:2), and PE(O-36:2)—were significantly elevated in mice fed the HCHF diet (Figure 5C). Mild food restriction led to further increases in these species; however, only PE(32:1) reached statistical significance.

Figure 5D highlights PE species previously reported to confer protective effects in CHD and CAD. Among these, PE(36:6) was significantly elevated in HCHF-fed mice subjected to mild food restriction, compared to chow-fed, food-restricted controls.

Phospholipid analysis of gallbladder bile is presented in supplementary Figure S4. Total PE levels did not differ significantly among the dietary groups. However, PE species bound to monounsaturated fatty acids (PE-MUFA) were significantly increased in mice on the HCHF diet. Unexpectedly, PE species bound to polyunsaturated fatty acids (PE-PUFA) remained unchanged. However, concentrations of several atherogenic PE species—PE(32:1), PE(34:1), PE(36:5), PE(38:2), and PE(O-36:2)—were significantly elevated in bile following HCHF feeding.

Additionally, we quantified lysophosphatidylcholine (LPC) levels in the livers of ApoE-deficient mice and observed a significant 33% reduction in total LPC content following the HCHF diet (Figure 5E). Consistent with trends seen in other phospholipid classes, there was a notable shift in LPC composition from PUFA-bound to MUFA-bound species (Figure 5F).

Regarding atherogenic LPC species, mice on the HCHF_DR diet showed a significant increase in LPC(16:1), while LPC(22:4) and LPC(22:5) were significantly decreased. In addition, several LPC species considered cardioprotective—namely LPC(16:0), LPC(18:0), LPC(18:2), and LPC(20:4)—were significantly reduced. Conversely, LPC(18:1) was significantly elevated (Figure 5H).

Taken together, although total hepatic LPC levels were significantly reduced by the HCHF diet, there was a selective enrichment of atherogenic LPC species, indicating a shift toward a more pro-atherogenic LPC profile.

In the bile, total LPC levels increased by more than 40% following mild food restriction, although this increase did not reach statistical significance (Supplementary Figure S5A). Similar to hepatic trends, LPC composition shifted markedly from PUFA-bound to MUFA-bound species (Supplementary Figure S5B). Furthermore, the atherogenic LPC(16:1) and the cardioprotective LPC(18:1) were significantly elevated in HCHF-fed animals, whereas LPC(18:2), another cardioprotective species, was significantly reduced (Supplementary Figure S5D and 5D).

### Quantification of clinically relevant pro-atherogenic sphingomyelin and ceramide acyl chains

We quantified total sphingomyelin (SM) and ceramide levels in liver tissue (Figure 6A). SM levels were significantly reduced in mice fed the HCHF diet, whereas mild food restriction restored hepatic SM concentrations to levels comparable to those in Chow-fed controls. Unlike other phospholipid classes, SM species bound to saturated fatty acids (SFA), monounsaturated fatty acids (MUFA), and polyunsaturated fatty acids (PUFA) were all significantly decreased following HCHF feeding. Importantly, these reductions were fully reversed by mild food restriction (Figure 6B).

**Figure 6.**
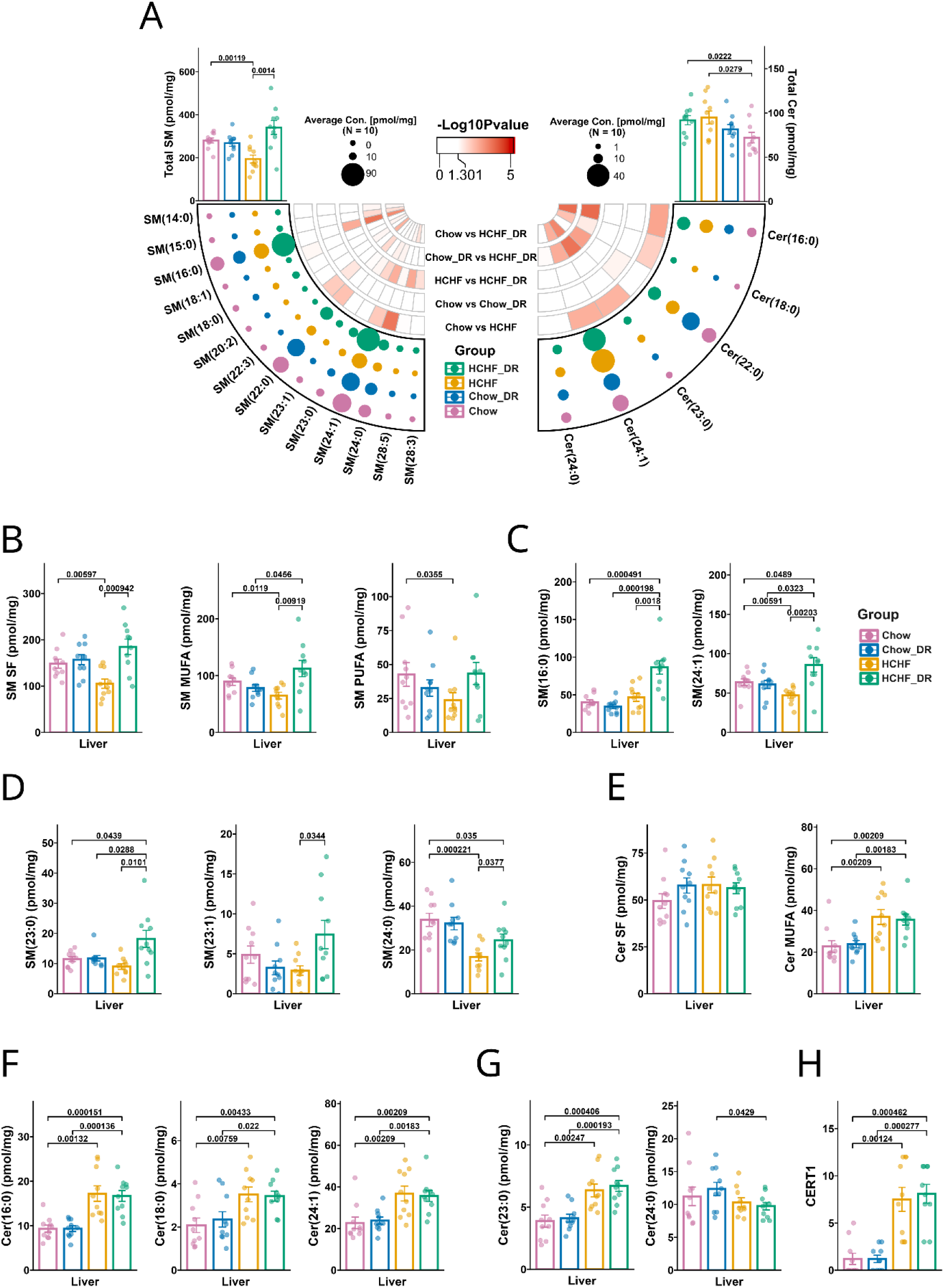
Quantification of hepatic sphingomyelin (SM) and ceramide (Cer) species. (A) Circular bubble heatmaps illustrating the composition and concentrations (pmol/mg tissue) of sphingomyelin species in the liver (left panel) and ceramides (right panel); red arcs denote -log₁₀ transformed P values for group comparisons. (B) Total hepatic sphingomyelin levels of saturated (SFA), monounsaturated (MUFA), and polyunsaturated (PUFA) SM species. (C) Quantification of pro-atherogenic SM species in the liver. (D) Quantification of anti-atherogenic SM species in the liver. (E) Total hepatic levels of Cer species grouped by saturation type. (F) Quantification of pro-atherogenic Cer species in the liver. (G) Quantification of anti-atherogenic Cer species in the liver. (H) Ceramide risk score of CERT1 in the liver. Data are presented as mean ± SEM. P values are shown where applicable.

Analysis of individual SM species showed that the atherogenic SM(16:0) and SM(24:1) were significantly elevated in HCHF-DR mice (Figure 6C). Additionally, mild food restriction led to significant increases in the cardioprotective SM(23:0) and SM(23:1), while SM(24:0), another cardioprotective species, was significantly decreased (Figure 6D).

Total ceramide levels increased by 30% following the HCHF diet, and this elevation was not significantly affected by mild food restriction (Figure 6A). Although mild food restriction in both Chow- and HCHF-fed animals was associated with an approximate 15% increase in ceramides bound to saturated fatty acids (SFAs), this change was not statistically significant. In contrast, the HCHF and HCHF-DR diets induced a statistically significant ∼38% increase in MUFA-bound ceramides. Additionally, the HCHF diet significantly elevated atherogenic ceramide species—Cer(16:0), Cer(18:0), and Cer(24:1)—with levels remaining unchanged by mild food restriction (Figure 6F). For cardioprotective ceramides, Cer(23:0) increased significantly, while Cer(24:0) decreased significantly after the HCHF diet; these effects remained stable by mild food restriction (Figure 6G).

Finally, we calculated the CERT1 ceramide risk score (Figure 6H), which revealed significantly elevated mean scores of 7.5 and 8.1 in ApoE mice fed HCHF and HCHF-DR diets, respectively. This modest increase in CERT1 aligns with the significant enlargement of atherosclerotic plaques observed following mild food restriction (Figure 1B).

Total sphingomyelin (SM) levels in bile were significantly increased in response to mild food restriction (Supplementary Figure S6A). Unlike the liver, SM species bound to saturated fatty acids (SFAs) were significantly elevated in the bile of ApoE mice fed the HCHF diet. Mild food restriction further enhanced MUFA-bound SM levels, with increases of 150% and 66% in the Chow-DR and HCHF-DR groups, respectively. Additionally, PUFA-bound SM was specifically elevated in the bile of Chow-DR animals (Supplementary Figure S6B). Note mild food restriction in HCHF-fed mice led to significant increases in atherogenic SM species, particularly SM(16:0) and SM(24:1) (Supplementary Figure S6C).

Total ceramide levels in bile were also significantly elevated following the HCHF diet (Supplementary Figure S6A). Furthermore, mild food restriction under the HCHF diet significantly increased MUFA-bound ceramides (Supplementary Figure S6D). When comparing dietary groups, atherogenic Cer(24:1) was significantly elevated in HCHF-fed mice relative to those subjected to mild food restriction (Supplementary Figure S6E). In contrast, the cardioprotective Cer(23:0) was significantly increased, while Cer(24:0) was significantly reduced in both Chow-DR and HCHF-fed animals. However, mild food restriction partially restored Cer(24:0) levels to those observed in Chow-fed controls (p = 0.0341; Supplementary Figure S6F).

Lastly, we calculated the bile-derived CERT1 ceramide risk score, which was significantly elevated in ApoE mice fed HCHF and HCHF-DR diets, with mean scores of 5.2 and 6.9, respectively.

## Discussion

Weight loss is generally recommended for patients with cardiovascular disease (CVD), particularly those who are overweight, as it can improve multiple metabolic risk factors ^1^. However, the impact of mild food restriction on lipid remodeling—especially with respect to pro-atherogenic lipid species—remains poorly understood. To address this, we employed ApoE-deficient mice, a well-established model for atherosclerosis that develops plaques resembling human lesions even on a chow diet, and shows accelerated plaque progression on a high-fat/Western diet. Using deep lipidomics, we identified clinically relevant atherogenic lipid species previously detected in nearly 500 CVD and 775 T2DM patients. By integrating these datasets through a consensus-driven approach, we provide evidence that our findings in ApoE-deficient mice have direct clinical relevance.

Specifically, the liver is a primary site for the synthesis of cholesterol, cholesterol esters (CE), and phospholipids, and clinical evidence demonstrates strong correlations between lipid profiles in the liver, adipose tissue, and plasma across diverse lipid classes and their acyl derivatives, reflecting a tightly coordinated metabolic network ^26^. In this study, we show that mild body weight loss in ApoE-deficient mice fed a high-fat diet led to significant increases in free cholesterol and pro-atherogenic phospholipids—including nine phosphatidylcholine (PC) species, six phosphatidylethanolamine (PE) species, one lysophosphatidylcholine (LPC), two sphingomyelins (SM), and three ceramides. Seven atherogenic CE species were also significantly elevated, whereas CE(20:0) did not reach statistical significance (p = 0.123). The rise in free cholesterol may be partly attributed to the marked repression of acyl-CoA:cholesterol acyltransferase 1 (ACAT1) under the HCHF diet, despite a mild, non-significant upregulation of ACAT2 (Supplementary Figure S2B). Importantly, Ces1f expression was suppressed more than tenfold. Given its established retinyl ester hydrolase activity and sequence homology to other cholesterol ester–hydrolyzing carboxylesterases, it is plausible that Ces1f also contributes to cholesterol ester hydrolysis, although this remains to be experimentally confirmed. Furthermore, hepatic lipase levels remained unchanged, suggesting it likely did not influence cholesterol release from lipoprotein surfaces in this context. Together, the repression of ACAT1 and Ces1f provides a plausible mechanistic explanation for the significant increase in free cholesterol despite unchanged total cholesterol (Figure 1C).

In stark contrast, mild weight loss induced by the same dietary restriction in mice on a Chow diet led to minimal changes in pro-atherogenic phospholipids, with only one species each of PC and LPC showing an increase. This dichotomy suggests that mild body weight loss under high-fat dietary conditions is uniquely associated with enhanced production of pro-atherogenic phospholipids, which correlated with a marked increase in plaque size (Figure 1).

Recent independent evidence demonstrates that caloric restriction can negatively impact atherosclerosis progression in young ApoE/LDL receptor knockout (ko) mice ^31^. While intermittent fasting on a Chow diet effectively resolved dyslipidemia and atherogenesis, it failed to reduce atherosclerosis in ApoE mice fed a HCHF diet ^32,33^. Similarly, time-restricted feeding reduced atherosclerosis in LDLR but not in ApoE mice mice given a HCHF diet ^34^, and alternate-day fasting likewise aggravated atherosclerosis in ApoE mice ^35^.

Our study highlights the complex interplay between diet, lipid metabolism, and atherogenesis, suggesting that mild weight loss under high-fat dietary conditions may paradoxically exacerbate lipid-driven pathways that promote atherosclerosis. We observed a significant increase in free cholesterol following mild caloric restriction of the HCHF diet, an effect not seen with the standard Chow diet. In contrast, total cholesterol levels in the Chow_DR group were reduced by 15.6%, although this change did not reach statistical significance. Elevated cholesterol levels are well-established drivers of foamy macrophage formation. Macrophages internalize LDL cholesterol, which contributes to plaque development by stimulating the secretion of inflammatory cytokines, enzymes, and growth factors to promote lipid accumulation, inflammation, and immune cell recruitment ^36,37^.

Consistent with our findings, independent studies have shown that alternate-day or intermittent fasting in ApoE-deficient mice elevates serum and plasma LDL cholesterol levels ^32,34,35^, further strengthening the connection between dietary interventions, cholesterol metabolism, and the progression of atherosclerosis.

Moreover, clinical studies have demonstrated that fasting increases serum total cholesterol and LDL cholesterol levels in healthy non-obese individuals ^38^ as well as in patients with anorexia nervosa ^39^. A medically supervised study involving 13 healthy Chinese participants who underwent a 10-day complete fast similarly reported a significant rise in LDL cholesterol^40^. Additionally, a recent systematic review highlighted the emerging phenomenon of intermittent fasting’s effects on hypercholesterolemia ^41^.

Given that plasma ceramides are predictive of cardiovascular death in patients with stable coronary artery disease and acute coronary syndromes ^23^, we calculated the cardiovascular event risk (CERT1) score. We observed a highly significant increase in the CERT1 score—7.5 and 8.1, respectively—in response to a HCHF diet administered either ad libitum or under mild food restriction. These elevated scores indicate a high probability of cardiovascular events and mortality, that in a clinical setting warrants aggressive management. Although the increase was not statistically significant, mild food restriction was associated with a slightly higher CERT1 score.

Deep lipidomic analyses in clinical studies have identified atherogenic cholesterol esters in plasma ^9^, while a newly developed multi-lipid score has been shown to predict health outcomes associated with specific dietary fat intake ^10^. In our study, we identified eight cholesterol esters with well-established pro-atherogenic properties (Figure 2), which were significantly elevated—ranging from 1.5-to 9-fold—following mild caloric restriction on a HCHF diet. A similar pattern was observed for phospholipids, with 18 pro-atherogenic species and three ceramides increasing by 1.5-to 5-fold.

Consistent with our findings, the prospective population-based Bruneck study identified CE(16:1) as significantly associated with cardiovascular disease (CVD) risk ^30^, a result independently validated in patients with T2DM and subclinical atherosclerosis ^28^. Furthermore, lipidomic analyses from subsets of the DAVIS trial (N=113) ^10^ and the EPIC-Potsdam cohort (N=1886 with T2DM and 1671 with CVD) provided strong evidence linking cholesterol esters, phospholipids, and ceramides with cardiovascular risk ^9^.

We integrated these insights using a consensus-driven approach and therefore propose that our findings in ApoE mice hold clinical relevance for identifying lipid-mediated pathways contributing to diet-induced atherogenesis.

### Study limitations

Several important caveats should be considered when interpreting findings from our study. First, fundamental differences in lipid metabolism between mice and humans limit the translatability of results. ApoE-deficient mice display an altered lipoprotein profile dominated by VLDL and chylomicron remnants, which contrasts with the LDL-centric profile typically observed in human atherogenesis. Second, plasma lipidomics and vascular tissue analysis will further strengthen the link between hepatic production of pro-atherogenic lipids and arteriosclerosis. Third, ApoE knockout mice develop spontaneous and accelerated atherosclerosis even on standard Chow diets, which may mask subtle effects of dietary interventions. This hyper-atherogenic phenotype may not accurately reflect the gradual disease progression characteristic of human CVD. Forth, the HCHF dietary interventions may exaggerate metabolic responses, limiting their real-world applicability. Fifth, ApoE-deficient mice generally lack the complex comorbidities common in humans—such as hypertension, diabetes, smoking exposure, and aging—that contribute to cardiovascular risk, thus failing to fully model CVD as a multifactorial disease process. Sixth, immunological differences represent another limitation. ApoE plays a role in immune regulation, and its absence alters inflammatory responses involved in atherosclerosis. Due to species-specific variations in immune function, findings related to inflammation and immune cell interactions may not fully translate to humans. Finally, our study used only male mice, thereby neglecting potential sex- and age-related differences in disease development and progression.

## Conclusion

In conclusion, our study reveals a paradoxical and potentially harmful effect of mild body weight reductions, which promotes the accumulation of pro-atherogenic cholesterol esters, phosphatidylcholines, and ceramide species, ultimately accelerating plaque growth and warrants a prospective clinical trial to ascertain the preclinical findings.

## Nonstandard Abbreviations and Acronyms

ACC/AHA: American College of Cardiology and American Heart Association
ADVANCE: The Action in Diabetes and Vascular Disease: Preterax and Diamicron-MR Controlled Evaluation
BA: Bile acids
BH: Benjamini–Hochberg
BMI: Body mass index
CARDIA: Coronary Artery Risk Development in Young Adults
CE: Cholesteryl esters
Cer: Ceramides
CERT1: Ceramide Test 1
CVD: Cardiovascular disease
cIMT: Carotid intima–media thickness
CLSA: The Canadian Longitudinal Study on Aging
DEGs: Differentially expressed genes
DR: Dietary restriction
ESI–MS/MS: Electrospray Ionization Tandem Mass Spectrometry
FC: Free cholesterol
FDR: False discovery rate
HCHF: High-cholesterol/high-fat
HexCer: Hexosylceramides
IS: Internal standards
LPC: Lysophosphatidylcholine
MRM: Multiple Reaction Monitoring
MUFA: Monounsaturated fatty acid
ORIGIN: ClinicalTrials.gov number NCT00069784, funded by Sanofi
PC: Phosphatidylcholine
PE: Phosphatidylethanolamine
PE(P): PE-based plasmalogens
PBS: Phosphate-buffered saline
PS: Phosphatidylserines
PUFA: Polyunsaturated fatty acid
RMA: Robust Multi-array Average
SFAs: Saturated fatty acids
SM: Sphingomyelins

## Acknowledgements

We would like to thank the EU and the German Ministry of Science and Education (BMBF) for financial support and Dr. R Schleef for donating tissue samples.

## Sources of Funding

This project was funded, in part, by the European Union LipidomicNet (‘Lipid droplets as dynamic organelles of fat deposition and release: translational research towards human disease’) The project was funded under the Seventh Framework Programme. Additionally, funding was obtained from the Virtual Liver Network (VLN) of the German Ministry of Science and Education (BMBF) [Grant number: 031 6154 to JB]. The funders had no role in study design, data collection and analysis, decision to publish, or preparation of the manuscript.

## Disclosures

The authors declare no conflict of interest.

## Supplemental Material

Tables S1-S6

Figures S1-S6

## Notes

### Competing Interest Statement

The authors have declared no competing interest.

